# Dynamics of centriole amplification in centrosome-depleted brain multiciliated progenitors

**DOI:** 10.1101/503730

**Authors:** Olivier Mercey, Adel Al Jord, Philippe Rostaing, Alexia Mahuzier, Aurélien Fortoul, Amélie-Rose Boudjema, Marion Faucourt, Nathalie Spassky, Alice Meunier

## Abstract

Centrioles are essential microtubule-based organelles organizing cilia and centrosomes. Their mode of biogenesis is semi-conservative: each pre-existing centriole scaffolds the formation of a new one, a process coordinated with the cell cycle. By contrast, multiciliated progenitors with two centrosomal centrioles massively amplify centrioles to support the nucleation of hundred of motile cilia and transport vital fluids. This occurs through cell type-specific organelles called deuterosomes, composed of centrosome-related elements, and is regulated by the cell cycle machinery. Deuterosome-dependent centriole amplification was proposed for decades to occur *de novo*, i.e. independently from pre-existing centrioles. Challenging this hypothesis, we recently reported an accumulation of procentriole and deuterosome precursors at the centrosomal daughter centriole during centriole amplification in brain multiciliated cells. Here we further investigate the relationship between the centrosome and the dynamic of centriole amplification by (i) characterizing the centrosome behavior during the centriole amplification dynamics and (ii) assessing the dynamics of amplification in centrosome-depleted cells. Surprisingly, although our data strengthen the centrosomal origin of amplified centrioles, we show limited consequences in deuterosome/centriole number when we deplete centrosomal centrioles. Interestingly, in absence of centrosomal centrioles, procentrioles are still amplified sequentially from a single focal region, characterized by microtubule convergence and pericentriolar material (PCM) self-assembly. The relevance of deuterosome association with the daughter centriole as well as the role of the PCM in the focal and sequential genesis of centrioles in absence of centrosomal centrioles are discussed.

## Introduction

Multiciliated cells (MCC) grow up to several hundred of motile cilia to generate fluid flow along organ lumen and promote essential respiratory, reproductive, and brain functions^1^. These cilia are nucleated by a basal body which is a mature centriole docked at the plasma membrane. Each multiciliated cell nucleates from 30 to 300 cilia, depending on the organ. Multiciliated precursors with only one centrosome must therefore amplify up to 300 centrioles to support cilia nucleation. This amplification occurs through the formation of intermediate organelles, the deuterosomes, which are composed of centrosome-related elements^2,3^. Deuterosomes support massive centriole production within a molecular cascade that is shared with the cell cycle centrosome duplication program^3–5^. Because two centrosomal centrioles seemed insufficient to scaffold the formation of tens of centrioles, massive centriole production through deuterosome structures was proposed to arise independently from the centrosome in MCC^6^. Challenging this postulate, an electron microscopy study showing association of deuterosomes with centrosomal centrioles in chick trachea proposed that the “procentriole clusters may form initially in close association with the diplosomal centrioles”^7^. More recently, we highlighted an atypical asymmetry between the mother and daughter centriole of the centrosome during cultured brain MCC differentiation, where deuterosomes are seen associated with the daughter centriole by electron microscopy^8^. Centriole amplification dynamics revealed by Cen2GFP live imaging together with centriole duplication players (Cep152, Plk4, Sas6) and Deup1 accumulation at the daughter centriole further suggested that procentrioles were amplified from the young centrosomal centriole through deuterosome formation. It also suggested the existence of a local micro-environment conductive to the formation of these auxiliary centrosome structures^8^. Interestingly, in mouse tracheal epithelial cell cultures, the transcription factor E2F4 was shown to accumulate in the centrosomal region during centriole amplification, and to be involved in deuterosome formation^9^. Here we further investigate the relationship between the centrosome organelle and the dynamics of centriole amplification by (i) characterizing the centrosome behavior during the centriole amplification dynamics and (ii) assessing the dynamics of amplification in cells depleted from one or both centrosomal centrioles.

## Results

### Centriole amplification proceeds sequentially and arises from the centrosomal region

We have recently shown that centriole amplification in mouse-brain progenitor cells is marked by 3 phases^8^. The amplification A-phase, during which procentrioles form around centrosome and deuterosome platforms, the growth G-phase, during which all procentrioles grow and mature synchronously from these platforms, and the disengagement D-phase, during which maturing centrioles detach from their growing platforms to dock apically, become basal bodies, and nucleate motile cilia. In order to precise A-phase dynamics, we used our *in vitro* assay where the differentiation of a mono-layered epithelium of ependymal cells from transgenic mice expressing a GFP-tagged version of the centriolar core protein Centrin 2 (Cen2GFP) allows the monitoring of centriole amplification dynamics (Fig 1a, b, supplementary movie 1)^8,10^. We first focused on the formation of immature procentrioles around deuterosomes (taking the form of Cen2GFP “halos”) during the A-phase and their subsequent maturation (procentriole polyglutamylation, widening, and lengthening^8^), visible by the transformation of Cen2GFP halos into “flowers” during the G-phase. We quantified that (i) the number of Cen2GFP halos increased over time in all the cells (Fig 1a, c; grey), suggesting that procentrioles are formed sequentially in the MCC progenitor, and (ii) the maturation of procentrioles at the A-to G-phase transition was timely correlated with a stop in the generation of new Cen2GFP structures (Fig 1a, c, d; orange), thus confirming that procentriole formation occurs only during A-phase.

**Figure 1:**
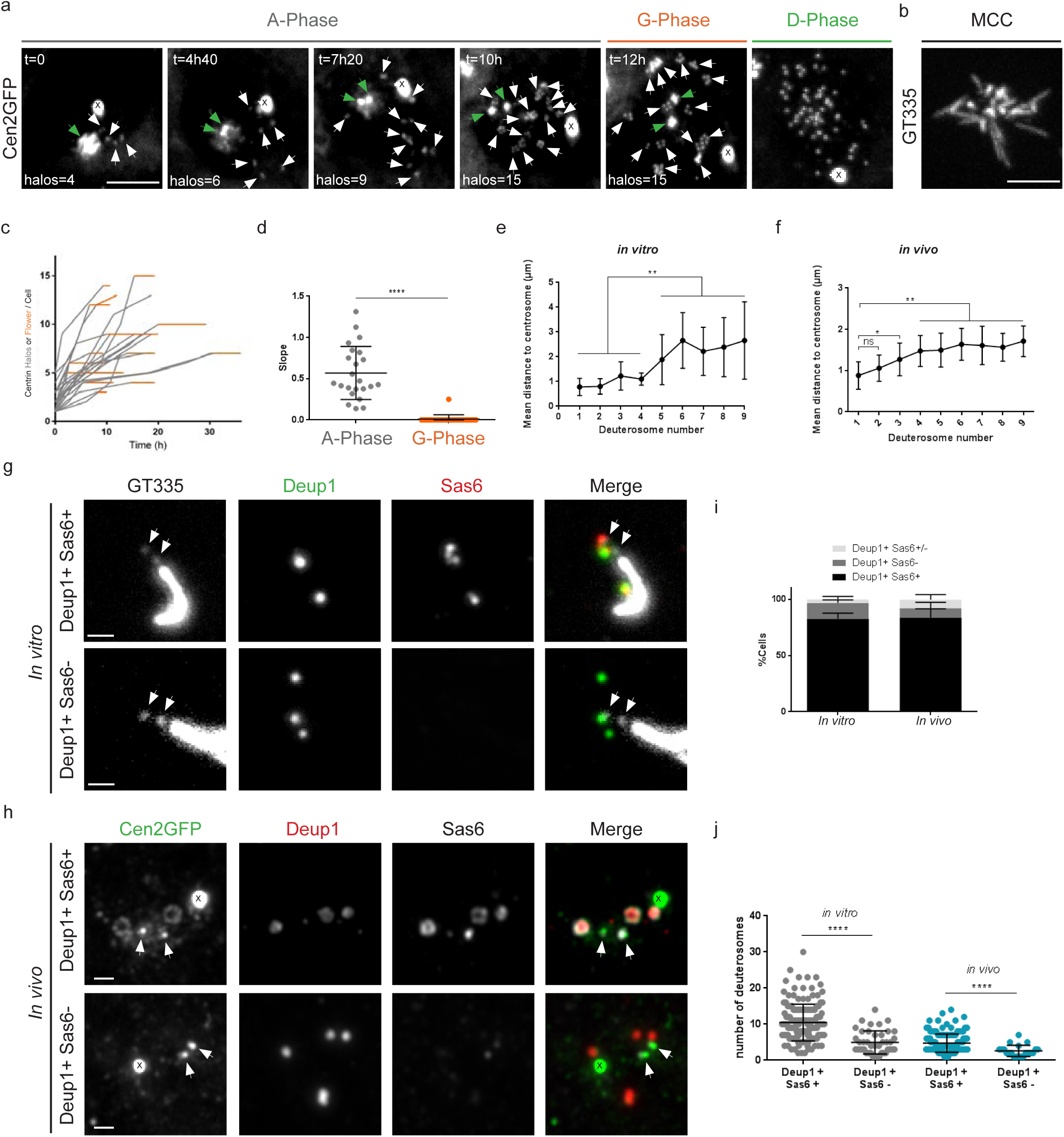
Centriole amplification proceeds sequentially and arises from the centrosomal region. **a**. Time lapse sequences of a Cen2GFP ependymal progenitor undergoing the different stages (A-Phase, G-Phase and D-Phase) of centriole amplification. Early A-Phase is characterized by Cen2GFP cloud surrounding centrosomal centrioles and the first visible halos in the nearby cytoplasm. As amplification progresses, the number of halos increases and they localize throughout the cytoplasm. In G-Phase, the final number of halos is reached, Cen2GFP halos transform into flowers where procentrioles are becoming visible. In D-Phase, procentrioles individualize. White arrows indicate centrin halos or flowers. Green arrows indicate centrosomal centrioles. **b**. Cilia immunostaining with GT335 antibody of a Cen2GFP ependymal MCC at the end of a time lapse experiment. **c**. Number of centrin halos (Gray) or flowers (Orange) during time lapse experiments in Cen2GFP ependymal progenitors (Δt=40min, n=23 cells). Timepoints are chosen when halos or flowers number is clearly visible. **d.** Trend line slopes corresponding to each cell observed in **c** in A-Phase and G-phase. **e, f**. Mean distances of deuterosomes to the centrosome depending on the number of deuterosomes in the cell *in vitro* (**e,** n=67 cells), and *in vivo* (**f,** n=134 cells). Deuterosomes were counted when positive for Deup1 and Sas6 stainings. **g and h**. Immunostaining of cells in A-phase with loaded (Deup1+/Sas6+) or unloaded (Deup1+/Sas6-) deuterosomes, *in vitro* (**g**), and *in vivo* (**h**). Arrows indicate centrosomal centrioles. **i.** Quantification of cells with loaded (Deup1+/Sas6+), partially loaded (Deup1+/Sas6+/-) or unloaded deuterosomes (Deup1+/Sas6-) *in vitro* (n=128 cells) and *in vivo* (n=279 cells). **j**. Quantification of the number of deuterosomes depending on the loading status of deuterosomes *in vitro* and *in vivo* (n=196 cells *in vitro,* n=165 cells *in vivo*). « X » indicates GFP aggregates. Scale bars: **a**,**b**: 5µm; **g**,**h**: 1µm.

In addition to the accumulation of centriole duplication players (Cep152, Plk4, Sas6) and Deup1 at the daughter centriole, the focal formation of the first Cen2GFP halos suggested that amplification initiates from the centrosomal region^8^ (Fig 1a, supplementary movie 1). Cloudy Cen2GFP accumulation around the centrosome at the beginning of centriole amplification however hindered accurate observation of the dynamic. To overcome this problem, we classified the cells depending on the number of deuterosomes they contain, using Deup1 and Sas6 immunostainings, assuming that early amplification would correspond to small number of deuterosomes. By measuring the distances between each Deup1/Sas6 positive deuterosome and the centrosome, we demonstrated that the first deuterosome was observed within a radius of 1µm from the centrosome both *in vitro* and *in vivo* (Fig 1e, f). Furthermore, the mean deuterosome-centrosome distance increased with the number of deuterosomes in the cell (Fig 1e, f). Altogether, this supports the scenario where centriole clusters organized around deuterosomes are sequentially formed within the centrosome prior to moving away in the cytoplasm^8^.

Because two studies report that Deup1 staining could be observed without procentriole markers^11,12^, we also decided to investigate whether this phenotype of “unloaded deuterosomes” is a reproducible step of differentiation in brain MCC. By carefully observing Deup1 and Sas6 stainings in cultured cells we found that around 20 % of cells in A-phase possessed Deup1 signal without Sas6 (Fig 1g, i). Because cells in culture can display abnormal features, we reproduced the experiments *in vivo,* in P2 mouse brain ventricles, and confirmed the *in vitro* result (Fig 1h, i). Although the mean number per cell of these “unloaded” deuterosomes was smaller than the mean number per cell of “loaded” deuterosomes, the distribution did not support the existence of a reproducible step of differentiation (Fig 1j). In fact, the majority of cells both *in vitro* and *in vivo* showed Sas6 loaded deuterosomes, even when very few deuterosomes are present (Fig 1i, j, Fig S4). This rather suggests that deuterosomes can appear unloaded in a subset of cells. Since the shape of the Deup1 signal of these “unloaded” deuterosomes is not always spherical or ring-shaped, correlative light and electron microscopy would be necessary to precise which Deup1 signal correlates to a bona fide electron dense deuterosome.

### An atypical centrosome cycle is observed during centriole amplification

Given the apparent centrosomal origin of amplified centrioles and our previous observation that Cep152, Plk4, Sas6 and Deup1 accumulates at the daughter centriole during A-phase^8^, we characterized centrosome behavior during the amplification process. Interestingly, we first found that the daughter centriole Cep164 staining evolved. The daughter centriole did not stain for Cep164 at the beginning of A-phase (Fig 2a). However, as the A-phase progressed and the number of deuterosome increased, the daughter centriole gradually stained positive for Cep164 to finally become indistinguishable from the mother centriole during the G-phase (Fig2a-c). This daughter-to-mother centriole conversion was confirmed by the growth of a second primary cilium during the G-phase (Fig 2d, e). Such bi-ciliated cells were also observed *in vivo* confirming that it is not an artifact of the culture system (Fig 2d). These two cilia then depolymerize during the disengagement phase, consistent with the migration of the centrosome and deuterosomes to the nuclear membrane^5^, and centrosomal centrioles are no longer distinguishable from mature disengaged procentrioles (Fig 2d, e). Using live imaging with high temporal resolution in cells with a limited number of procentrioles, we were able to track centrosomal centrioles and showed that they remained apically localized during procentriole migration. They were eventually joined by the procentrioles to form the basal body patch. This was confirmed by fate-tracing centrosomal centrioles using Cen2GFP/Cen-RFP pulse-chase experiments^13^. Altogether, these data highlight an atypical centrosome cycle, correlated in space and time to the progression of procentriole amplification in brain MCC. Most interestingly in this cycle, (i) the sequential formation of procentriole clusters during A-phase arises from the centrosomal region and (ii) the arrest in cluster formation, marked by procentriole growth at the A-to G-transition, correlates with the maturation of the centrosomal daughter centriole (Fig 2f).

**Figure 2:**
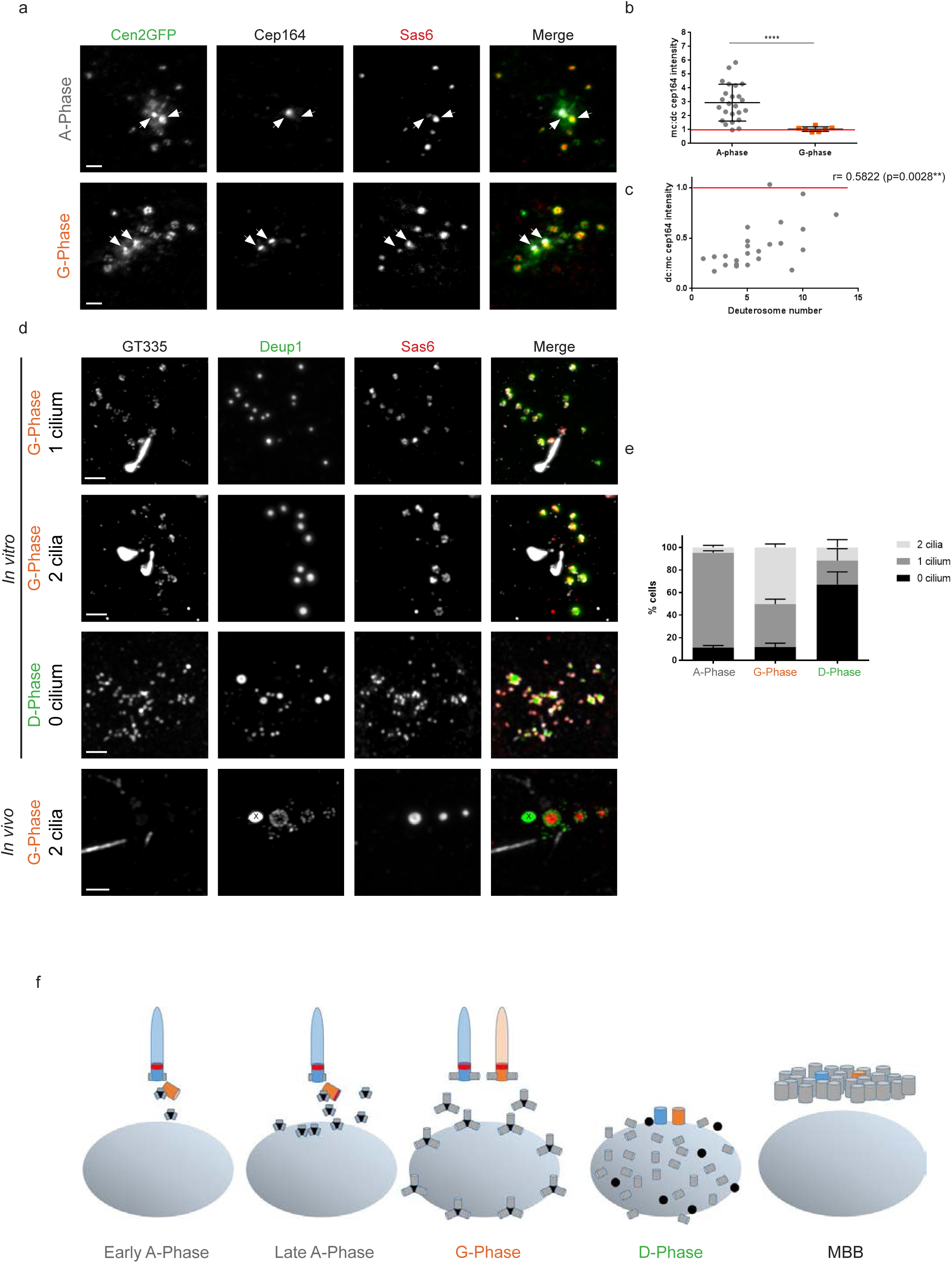
An atypical centrosome cycle is observed during centriole amplification. **a**. Cep164 and Sas6 immunostainings on Cen2GFP ependymal progenitor during A- and G-Phases. Arrows indicate centrosomal centrioles. **b**. mother:daughter centriole (mc:dc) Cep164 signal ratios in A-phase (n=24 cells) and G-Phase (n=7 cells). **c**. Correlation between mc:dc Cep164 signal ratios and deuterosome numbers in the cells during A-Phase (n=24 cells from **b**). **d**. GT335, Deup1 and Sas6 immunostaining on Cen2GFP ependymal progenitor during G- and D-Phase *in vitro* and G-phase *in vivo*. **e**. Percentage of cells with 0, 1 or 2 GT335 positive cilia in A-, G-and D-phases *in vitro* (n=210, 73 and 21 cells in A-, G- and D-phase respectively). **f**. Representative scheme of centrosomal centriole behaviour during centriole amplification. Nucleus is represented in gray, deuterosomes in black, procentrioles in gray, Cep164 in red, mother centriole in blue, daughter centriole in orange. « X » indicates Cen2GFP aggregate. Scale bars, 2µm.

### Depletion of centrosomal centrioles slightly increases centriole amplification in brain MCC

Given the tight spatio-temporal link between centrosome behavior and centriole amplification shown here and previously^8^, we wanted to evaluate the role of the centrosomal centrioles in the process. We used the small molecule inhibitor centrinone^14^, which inhibits Plk4 kinase function, to deplete centrosomal centrioles from progenitor cells before triggering MCC differentiation (Fig 3a). By treating primary progenitor cells with centrinone during the proliferating phase, we obtained a mixed population of cells with 2 centrosomal centrioles (2cc), 1 centrosomal centriole (1cc) or 0 centrosomal centriole (0cc) (Fig 3a,b). Although centrinone-treated proliferating progenitors reached confluence later when compared to control (not shown), they were able to proliferate confirming that mouse cells are less sensitive than human cells to centrosome-loss for cell division^6,14–18^. MCC differentiation was then triggered after centrinone wash out and high confluence cell seeding in serum free medium. We obtained heterogeneous cultures with equivalent proportions of 2cc, 1cc and 0cc cells. These proportions did not vary between the first day of differentiation (Day *In Vitro* 0 or DIV0) and DIV5 suggesting that centriole-depleted cells are not significantly more prone to cell death (Fig 3c). Using single-cell approaches, we first confirmed by correlative light and electron microscopy (CLEM) that Cen2GFP was suitable to assess the presence of centrosomal centrioles in the cells (Fig S1, S2). Then, using immunostainings and CLEM, we sought to characterize centriole amplification in the 3 cell populations. Interestingly, parental centriole depletion neither blocked the MCC progenitor capacity to form canonical deuterosomes nor to amplify procentrioles (Fig 3d, e; Fig S1, S2). However, 0cc cells possessed slightly more deuterosomes than 1cc or 2cc cells, suggesting that centrosomal centrioles could tune this parameter (Fig 3f). As previously shown^8^, Deup1 accumulation was observed at the daughter centriole in 2cc cells. Interestingly, removing the daughter centriole (1cc cells) did not change the incapacity of the mother centriole to accumulate Deup1 (Fig 3g), confirming an asymmetry in the propensity of the mother versus daughter micro-environment for Deup1 concentration and putative deuterosome formation. Next, we scored the dynamics of amplification using single cell live imaging and found that the 3 populations achieved the stereotypical phases of centriole amplification (A-, G- and D-phases) with a similar spatio-temporal pattern (Fig 3h, i). Consistent with the increase in deuterosome number, a slight increase in the number of centrioles was observed in 0cc cells as compared to 2cc cells (Fig 3j). Altogether our data suggest that in multiciliated cells like in cycling cells, resident centrioles are dispensable organelles for centriole biogenesis, which might participate in the deuterosome/centriole copy number control. Whether the structure of procentrioles is affected by the absence of parental centrioles needs further investigation^19^.

**Figure 3:**
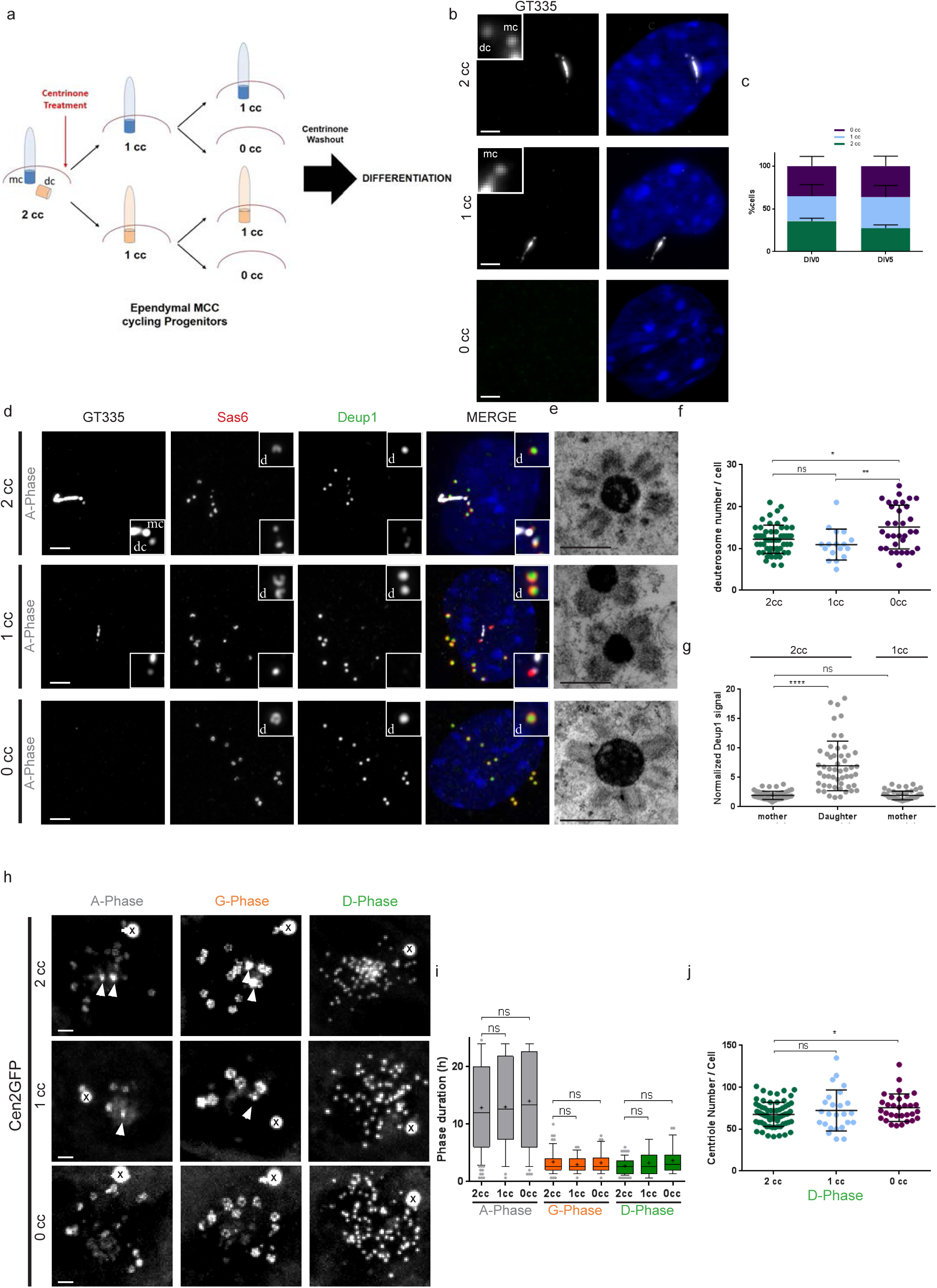
Depletion of centrosomal centrioles slightly increases centriole amplification in brain MCC. **a**. Experimental procedure used to deplete centrosomal centrioles in MCC cycling progenitors using centrinone. See methods. 2cc, 1cc or 0cc for 2, 1 or 0 centrosomal centriole, respectively. **b**. Representative pictures of 2cc, 1cc or 0cc ependymal progenitor cells stained with GT335. **c**. Repartition of 2cc, 1cc or 0cc cell populations at two different days of differentiation (DIV for Day *In Vitro*). Cells selected for this quantification are ependymal progenitors that have not started centriole amplification (« centrosome » stage; n=912 and 566 cells at DIV0 and DIV5 respectively). **d**. Representative GT335, Deup1 and Sas6 immunostainings of 2cc, 1cc or 0cc cells during A-Phase. Zoom in pictures highlight deuterosomes « d » and centrosomal centrioles (mc: mother centriole and dc: daughter centriole). **e**. Representative EM pictures of deuterosomes in 2cc, 1cc or 0cc cells. The status of the centrosome was identified by correlative light and electron microscopy. **f**. Quantification of deuterosome number per cell in 2cc, 1cc or 0cc cells during G-Phase (n= 55, 17, 33 cells for 2cc, 1cc and 0 cc respectively). **g**. Normalized Deup1 signal (centrosomal centriole:cytoplasmic Deup1 signal) on mother and daughter centriole in 2cc cells, or on mother centriole in 1cc cells during A-Phase (n = 52 and 54 cells for 2cc and 1cc respectively). **h**. Sequences from time lapse experiments on 2cc, 1cc or 0cc Cen2GFP cells. Note that a Cen2GFP cloud is still present in 0cc cells. **i**. Box (25 to 75%) and whisker (10 to 90%) plots of A-(Gray), G- (Orange), and D- (Green) phase duration in 2cc, 1cc or 0cc Cen2GFP progenitors. Lines indicate medians, and crosses indicate means. 2cc cells have been taken from DMSO and Centrinone treated cultures (In A-Phase: n=107 cells for 2cc; n=36 cells for 1cc; n=49 cells for 0cc; In G-Phase: n=73 cells for 2cc; n=32 cells for 1cc; n=36 cells for 0cc; In D-Phase: n=65 cells for 2cc; n=25 cells for 1cc; n=28 cells for 0cc). **j**. Final centriole number counted during D-Phase in 2cc, 1cc or 0cc Cen2GFP progenitors. Because centrosomal centrioles are no longer distinguishable from maturing procentrioles in D-Phase, quantification is performed after Cen2GFP videomicroscopy to know the orignal centrosome status of the cells. 2cc cells have been taken from DMSO and Centrinone treated cultures (n=69 cells for 2cc; n=25 cells for 1cc; n=31 cells for 0cc). Arrows indicate centrosomal centrioles. « X » indicates GFP aggregates. Scale bars, 2µm for optical microscopy, 500nm for EM.

### Centrosomal centriole-depleted cells form procentrioles within an acentriolar PCM cloud

Finally, we focused on 0cc cells in order to decipher the precise dynamics of *de novo* centriole amplification. *De novo* centriole assembly has been induced in human cycling cells depleted from centrosomal centrioles using laser ablation^20,21^, centrinone treatment^14^ or an auxin-dependent Plk4 degradation system^18^. The outnumbered centrioles were proposed to arise all at once during S-phase^21^ either from a PCM cloud^20^ or stochastically throughout the cytosol^18^. Using live imaging on 0cc MCC progenitors, we observed that Cen2GFP halos assemble sequentially, within a normal A-phase duration (Fig 3i), at a similar rate when compared to 2cc cells (Fig 4a, c, d, supplementary movie 2), and give rise to basal bodies growing cilia (Fig 4b). Since the sequential formation of halos was proposed to be due to their sequential generation from the daughter centriole^8^, we then analyzed their origin in absence of centrosomal centrioles. Resident centriole depletion by centrinone in human interphase cells has been shown to involve microtubule organization from the Golgi apparatus or from multiple cytoplasmic foci^22,23^. In MCC progenitors, the existence of a focal Cen2GFP cloud, from where the halos arose (Fig 4a, supplementary movie 2), suggested that a single acentriolar MTOC was self-organizing in the absence of resident centrioles. Consistently, the Cen2GFP cloud localized at the center of convergence of the microtubule network (Fig 4e, Fig S3a) and colocalized with a single Pericentrin/Cdk5Rap2 cloud (Fig 4f, Fig S3b, c). This acentriolar MTOC did not appear connected to the Golgi apparatus (Fig S3d). Interestingly, while the Pericentrin cloud surface was not modified by the status of the centrosome (Fig 4g), its size correlated with the number of deuterosomes in both 0cc and 2cc cells (Fig 4h). By measuring the mean distance of deuterosomes to the PCM cloud center, we revealed comparable results for 2cc and 0cc cells where the mean distance of deuterosomes to the PCM center increased with the number of deuterosomes (Fig 4i). Altogether, this suggests that, in the presence or absence of centrosomal centrioles, deuterosomes and their procentrioles arise sequentially from a single focal region, characterized by microtubule convergence and PCM self-assembly (Fig 4j).

**Figure 4:**
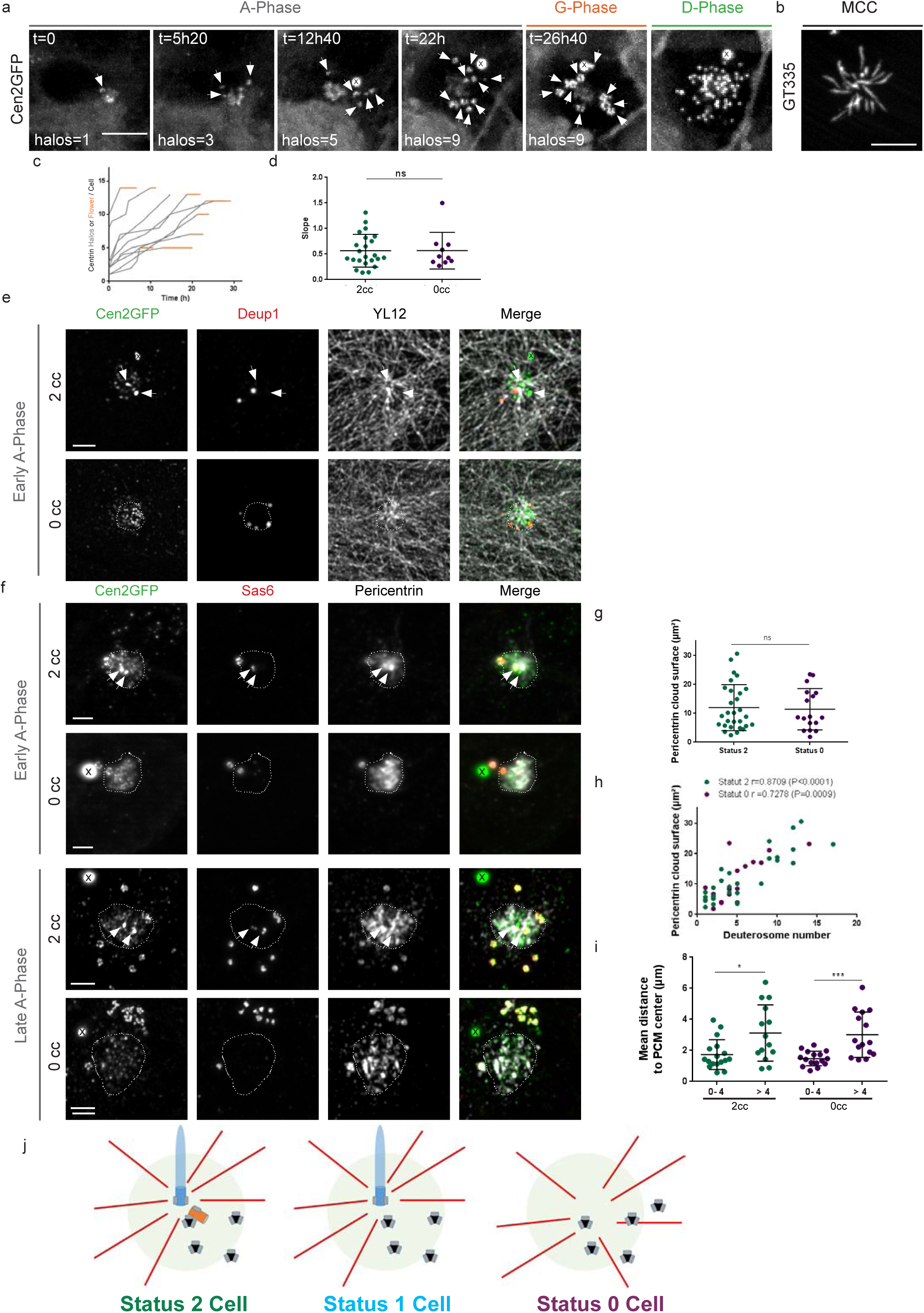
Centrosomal centriole-depleted cells form procentrioles within an acentriolar PCM cloud. **a**. Live imaging of a Cen2GFP ependymal progenitor depleted from centrosomal centrioles undergoing the different stages (A-, G- and D-Phase) of centriole amplification. **b**. GT335 immunostaining of a 0cc Cen2GFP ependymal MCC at the end of the time lapse experiment. **c**. Number of centrin halos (Gray) or flowers (Orange) during time lapse experiments in 0cc Cen2GFP ependymal progenitors (Δt=40min, n=10 cells). Timepoints are chosen when halos or flowers number is clearly visible. **d**. Comparison of centrin halo formation rate between 2cc or 0cc cells during A-Phase. Each dot represents the trendline slope corresponding to a cell observed during A-Phase in Fig 1c (2cc cells) and Fig 4c (0cc cells). **e**. Immunostainings showing deuterosome localization (Deup1) and the tyrosinated tubulin network (YL12) in 2cc or 0cc cells during early A-Phase. In 2cc cells, microtubule network converges on one centrosomal centriole whereas in 0cc cells, it converges on Cen2GFP cloud. Dashed line delineates Cen2GFP cloud. **f**. Representative pictures of Pericentrin and Sas6 immunostainings during early or late A-Phase in 2cc or 0cc cells. Dashed line delineates the Pericentrin cloud. **g**. Surface of the Pericentrin cloud in 2cc and 0cc cells during A-phase (n=29 cells for 2cc, n=17 cells for 0cc). **h.** Correlation between the Pericentrin cloud surface and the number of deuterosome in the cell in 2cc (green) or 0cc (purple) cells (n=29 cells for 2cc, n=17 cells for 0cc). **i**. Mean distance of deuterosomes to PCM center depending on the number of deuterosomes in the cells for 2cc and 0cc cells (for 2cc: 0-4 deuterosomes: n=17 cells; > 4 deuterosomes: n=14 cells. for 0cc: 0-4 deuterosomes: n=16 cells; > 4 deuterosomes: n=15). **j**. Model for centriole amplification in 2cc, 1cc or 0cc cells. PCM cloud is represented in light green, deuterosomes in black, procentrioles in gray, microtubules in red. Arrows indicate centrosomal centrioles. « X » indicates GFP aggregates. Scale bars: **a**, **b**: 5µm: **e**, **f**: 2µm.

## Discussion

In this study, we highlighted an atypical centrosome cycle, correlated in space and time to the progression of procentriole amplification in brain MCC. Most interestingly in this cycle, (i) the sequential formation of procentriole clusters during A-phase arises from the centrosomal region and (ii) the arrest in cluster formation, marked by procentriole growth at the A- to G-transition, correlates with the maturation of the centrosomal daughter centriole. Concurrent maturation of the young centrosomal centriole and the procentrioles is also seen in cycling cells and was shown to be Plk1 dependent^24^. Since Plk1 regulates the A- to G-transition in brain MCC progenitors^5^, it is tempting to speculate that it also drives the concurrent maturation of the two generations of centrioles in these differentiating cells. Interestingly, such maturation of the daughter centriole is also observed in MCC induced from primary fibroblasts^11^. The existence of a causal link between daughter centriole maturation and/or procentriole growth with the amplification arrest remains to be determined.

Although our data strengthen the centrosomal origin of amplified centrioles, cells depleted from centrosomal centrioles only show a minor defect in the production of deuterosomes and procentrioles. Comparable to what occurs in cycling cells, centrosomal centrioles, while dispensable, seem to restrict deuterosome and centriole number. This is however very limited relative to the final number of deuterosomes and centrioles produced in MCC. The fact that 1cc cells, where the mother centriole is still present, and 2cc cells show no difference suggests that that microtubule or PCM organization could be the parameters influencing deuterosome/centriole number. In this line, the “*de novo*” centriole formation in 0cc cells seems to occur within a PCM cloud that self-assembles in the absence of centrosomal centrioles.

Accumulation of centriole duplication players at the daughter centriole during A-phase, together with the observed association of deuterosomes with the proximal part of the daughter centriole^8^, suggests that, similar to cycling cells, a centriolar wall may serve as a preferential but dispensable nucleation site ensuring a structural patterning^19^ and/or facilitating nucleation events^25,26^. Whether the structure of centrioles is affected by the absence of parental centriole needs further investigation^19^. We can’t exclude the possibility that only a subset of deuterosomes/procentrioles is nucleated on the daughter centriole, while the others are formed in close proximity within the PCM. Or even that all deuterosomes are formed within the PCM and some only stick to the daughter centriole from the beginning to the end of A-phase biasing our interpretation. *In vivo*, tight proximity of Deup1 structures with the daughter centriole is verified but less systematic than *in vitro* and than what could have been expected by Sas6 asymmetry *in vivo*^8^ (Fig S4). Together with the existence of “unloaded deuterosomes” in the cytoplasm in a subset of cells, it suggests that deuterosome and procentriole nucleation can be uncoupled. Super-resolution live imaging on cells from Deup1-tagged knock-in mice with physiological concentrations of Deup1 may help to precise the dynamics of deuterosome formation in relation to the daughter centriole over-seeding capacity and to procentriole amplification in general.

Interestingly, even in absence of the daughter centriole (1cc cells), the mother centriole seems unable to accumulate Deup1. Since the PCM seems to be the site for deuterosome and procentriole assembly in the absence of centrosomal centrioles, mother/daughter asymmetry may reflect the existence of sub-compartments within the PCM that would be more prone to deuterosome and/or procentriole nucleation events to occur. In this line, increasing evidence suggests that the centriolar matrix, and particularly Pericentrin, regulates centriole assembly, and not only the other way around. Overexpression of Pericentrin leads to overduplication of centrioles in human transformed cells^27^. More recently, Pericentrin was shown to recruit Sas6 and to be involved in centriole biogenesis and stability in drosophila^28,29^. The increasing expression of Pericentrin during A-phase, visible by the increasing surface of the Pericentrin cloud, may be involved in the sequential formation of procentrioles we observed, even in the absence of centrosomal centrioles. Depleting pericentrin and driving the daughter centriole away from the mother centriole^30^ may allow to test the association of PCM with the daughter centriole and their respective contribution in MCC centriole amplification.

## Acknowledgements

We thank all members of the Spassky laboratory for comments and discussions. We thank K. Oegema and A.K. Shiau (Ludwig Institute for Cancer Research, La Jolla, CA) for sharing the Centrinone, E.A. Nigg for the Cep164 antibody; A.-K. Konate and R. Nagalingum for administrative support; the IBENS Animal Facility for animal care. We thank the IBENS Imaging Facility, with grants from Région Ile-de-France (NERF 2011-45), Fondation pour la Recherche Médicale (FRM) (DGE 20111123023), and Fédération pour la Recherche sur le Cerveau–Rotary International France (2011). The IBENS Imaging Facility and the team received support from Agence Nationale de la Recherche (ANR) Investissements d’Avenir (ANR-10-LABX-54 MEMO LIFE, ANR-11-IDEX-0001-02 PSL* Research University). The laboratory is supported by INSERM, CNRS, École Normale Supérieure (ENS), ANR (ANRJC JC-15-CE13-0005-01), European Research Council (ERC Consolidator grant 647466), FRM (Equipe FRM20140329547), Cancéropôle Ile-de-France (2014-1-PL BIO-11-INSERM 12–1). The authors declare no competing financial interests.

## Author contributions

O.M. designed the study, performed, analyzed experiments and wrote the manuscript; A.A.J. contributed to characterizing centrosome behavior during centriole amplification. P.R. contributed to correlative light and electron microscopy; N.S, A.Ma., A.F. and A-R.B. contributed to *in vivo* experiments; M.F. contributed to cell culture experiments; N.S. analyzed data, contributed to scientific discussions and edited the manuscript; A.M. initiated the study, supervised the project and wrote the manuscript.

## Data availability

All data generated or analysed during this study are included in this published article (and its Supplementary Information files).

## Methods Animals

All animal studies were performed in accordance with the guidelines of the European Community and French Ministry of Agriculture and were approved by the Direction départementale de la protection des populations de Paris (Approval number APAFIS#9343-201702211706561 v7). The mice used in this study have already been described and include: OF1 (Oncins France 1, Charles River Laboratories); Cen2GFP (CB6-Tg(CAG-EGFP/CETN2)3-4Jgg/J, The Jackson Laboratory). Videomicroscopy experiments were performed on cells from homozygous Cen2GFP mice. Other experiments were performed in parallel using OF1 or homozygous Cen2GFP mice; no differences between these two strains were observed regarding ependymal differentiation, amplification stages, centrosome asymmetry or number of deuterosomes and centrioles.

### Brain dissections

Whole mounts of developing ventricular walls were prepared from P2-P4 adult mice as previously described ^1^.

### Primary ependymal cell cultures and centrinone treatment

Newborn mice (P0–P2) were killed by decapitation. The brains were dissected in Hank’s solution (10% HBSS, 5% HEPES, 5% sodium bicarbonate, 1% penicillin/streptomycin (P/S) in pure water) and the extracted ventricular walls were cut manually into pieces, followed by enzymatic digestion (DMEM glutamax, 33% papain (Worthington 3126), 17% DNase at 10 mg ml^−1^, 42% cysteine at 12 mg ml^−1^) for 45 min at 37 °C in a humidified 5% CO2 incubator. Digestion was stopped by addition of a solution of trypsin inhibitors (Leibovitz Medium L15, 10% ovomucoid at 1 mg ml^−1^, 2% DNase at 10 mg ml^−1^). The cells were then washed in L15 and resuspended in DMEM glutamax supplemented with 10% fetal bovine serum (FBS) and 1% P/S in a Poly-l-lysine (PLL)-coated flask. Ependymal progenitors proliferated for 5 days until confluence followed by shaking (250 rpm) overnight. Pure confluent astroglial monolayers were replated at a density of 7 × 10^4^ cells per cm^2^ (corresponding to days *in vitro* (DIV) −1) in DMEM glutamax, 10% FBS, 1% P/S on PLL-coated coverslides for immunocytochemistry experiments, Lab-Tek chambered coverglasses (Thermo Fisher Scientific) for time-lapse experiments or glass-bottomed dishes with imprinted 50 µm relocation grids (Ibidi, catalogue no. 81148, Biovalley) for correlative Light/EM and maintained overnight. The medium was then replaced by serum-free DMEM glutamax 1% P/S, to trigger ependymal differentiation gradually *in vitro* (DIV 0). Centrinone was added on day2 of the proliferation phase at a final concentration of 0.6 µM. Centrinone was washed out 3 times in PBS on day5 of the proliferation phase just before trypsinisation and replating at high confluence for MCC differenciation.

### Immunostainings

Lateral brain ventricles were first pre-permeabilized in 0.2% Triton X-100 BRB medium (80mM PIPES, 1mM MgCl2, 1mM EGTA) for 2 min before fixation. Tissues or cell cultures (between DIV2 and DIV6) were then fixed in methanol at −20 °C for 10 min. Samples were pre-blocked in 1× PBS with 0.2% Triton X-100 and 10% FBS before incubation with primary and secondary antibodies. Tissues or cells were counterstained with DAPI (10 μg ml-1, Sigma) and mounted in Fluoromount (Southern Biotech). The following antibodies were used: rabbit anti-Cep164 (1:750) ^2^; rabbit anti-Deup1 (1:2000) (homemade, raised against the peptide TKLKQSRHI); mouse IgG1 anti-GT335 (1:500, Adipogen); mouse IgG2b anti-Sas6 (1:750, Santa Cruz); rabbit anti-Pericentrin (1:2000); rat anti-YL12 (1:500) and species-specific Alexa Fluor secondary antibodies (1:400, Invitrogen).

### Microscopy

#### Epifluorescence microscopy

Fixed cells and whole-mount ventricles were examined with an upright epifluorescence microscope (Zeiss Axio Observer.Z1) equipped with an Apochromat ×63 (NA 1.4) oil-immersion objective and a Zeiss Apotome with an H/D grid. Images were acquired using Zen with 240-nm z-steps.

#### Confocal super-resolution microscopy

Confocal image stacks were collected with a 63 x/1.4 Oil objective on an inverted LSM 880 Airyscan Zeiss microscope with 440, 515, 560 and 633 laser lines.

#### Videomicroscopy

Cultured cells between DIV2 and DIV6 were filmed *in vitro* using an inverted spinning disk Nikon Ti PFS microscope equipped with oil-immersion ×63 (NA 1.32) and ×100 (NA 1.4) objectives, an Evolve EMCCD Camera (Photometrics), dpss laser (491 nm, 25% intensity, 70-100ms exposition), appropriate filter sets for DAPI/FITC/TRITC, a motorized scanning deck and an incubation chamber (37 °C; 5% CO2; 80% humidity). Images were acquired with Metamorph Nx with 40 min time intervals. Image stacks were recorded with a z-step of 0.7 µm. After film acquisition, cells were then fixed for 5 min with 0.5% paraformaldehyde (PFA) using a Pasteur pipette without moving the chambered coverglass, and GT335 primary and secondary antibodies were added together for 25 min, in medium supplemented with FBS (10%) before the final images were acquired.

#### Correlative light and electron microscopy (CLEM)

Primary Cen2GFP ependymal progenitors were grown in 0.17-mm thick glass dishes with imprinted 50 µm relocation grids (Ibidi). At 3-6 days in vitro (DIV 3-6), cells were fixed with 4% PFA for 10 min and ependymal progenitors undergoing A-phase or G-Phase were imaged for Cen2GFP and DAPI, in PBS, with upright epifluorescence microscope (Zeiss Axio Observer.Z1). Coordinates on the relocation grid of the cells of interest were recorded. Cells were then treated for transmission electron microscopy. Briefly, culture cells were treated with 1% OsO4, washed and progressively dehydrated. The samples were then incubated in 1% uranyl acetate in 70% methanol, before final dehydration, pre-impregnation with ethanol/epon (2/1, 1/1, 1/2) and impregnation with epon resin. After mounting in epon blocks for 48 h at 60 °C to ensure polymerization, resin blocks were detached from the glass dish by several baths in liquid nitrogen. Using the grid pattern imprinted in the resin, 50 serial ultra-thin 70-nm sections of the squares of interest were cut on an ultramicrotome (Ultracut E, Leica) and transferred onto formvar-coated EM grids (0.4 × 2 mm slot). The central position of the square of interest and DAPI staining were used to relocate and image the cell of interest using a Philips Technai 12 transmission electron microscope.

### Quantification and statistics

After live imaging, rates of Cen2GFP structures appearance were measured. We quantified the number of centrin halos or flowers at different times and computed the slopes given by a linear regression. Distances of deuterosomes to the centrosome were measured between the deuterosome center of mass and the middle of the segment drawn between the two centrosomal centrioles. The mean distance was then calculated within each single cell. For the mean distance of deuterosomes to Pericentrin cloud center, region of interest corresponding to the Pericentrin cloud was manually drawn and centroid coordinates were then used as the PCM center. The same protocol was used for Pericentrin cloud surface measurements. Fluorescence signal intensities were quantified using Image J. All graphs and statistical analyses were obtained using GraphPad Prism software. Data were obtained from at least three independent experiments and the results presented as the mean ± s.d. Non-parametric two-tailed Mann–Whitney U-tests were used to compare groups of data. Pearson’s correlation coefficient was calculated to determine the strength of the relationship between two variables. For P-Values: ns: P>0.05; *: P≤0.05; **: P≤0.01; *** P≤0.001; **** P≤0.0001

---

**Supplementary Figure 1:**
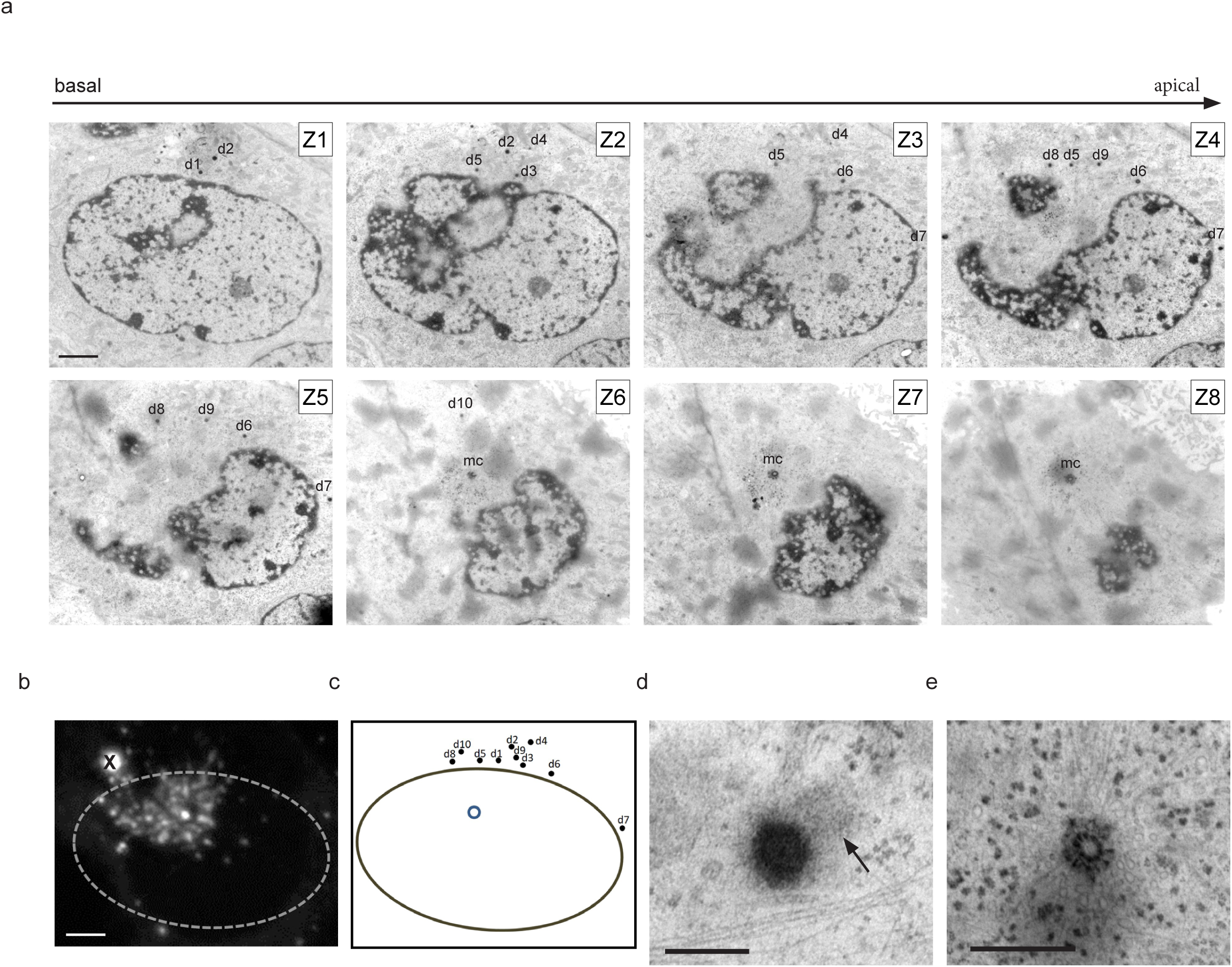
Characterization of a daughter centriole-depleted cell. **a**. EM serial pictures of a daughter centriole-depleted cell (1cc). 1cc cell is first selected using Cen2GFP signal (**b**). First representative picture (Z1) corresponds to the first ultrathin section (70nm) where deuterosomes « d » are observed. Deuterosomes are then counted in order of appearance. mc: mother centriole. Scalebar = 2µm. **b**. Cen2GFP signal of the daughter centriole-depleted cell observed in EM. Dashed line represents the nucleus. « X » indicates Cen2GFP aggregate. Scale bar = 2µm. **c**. Schematic representation of the cell observed in CLEM. Black dots represents deuterosomes, Solid line indicates nucleus and blue circle represents mother centriole. **d**. Zoom-in of a deuterosome with one growing procentriole. Arrow indicates the procentriole. Scale bar = 200nm. **e**. Zoom-in on the mother centriole surrounded by aggregates and organizing microtubules. Scalebar = 1µm

**Supplementary Figure 2:**
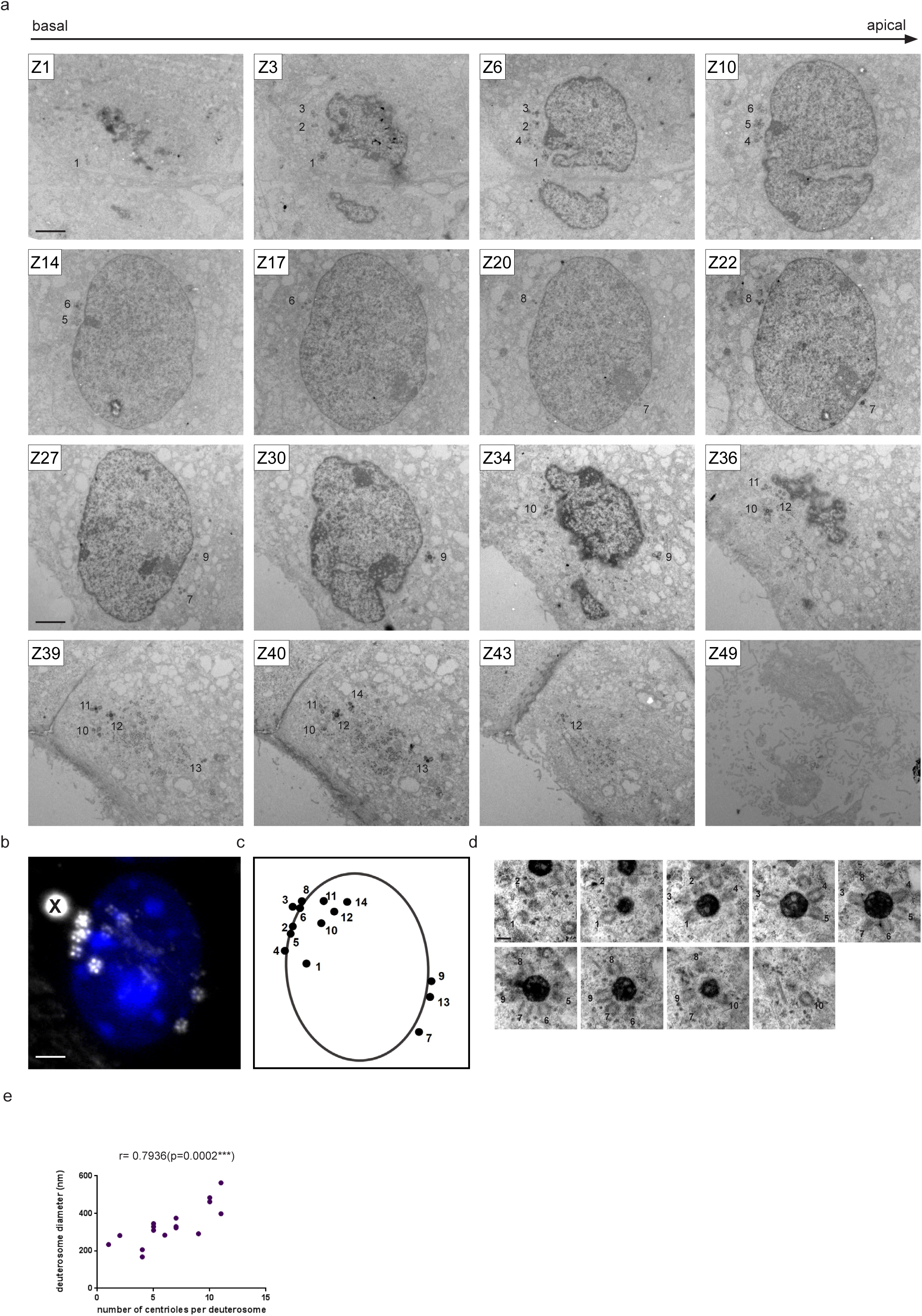
Characterization of a centrosomal centriole-depleted cell. **a**. EM serial pictures of a parental centriole-depleted cell (0cc). 0cc cell is first selected using Cen2GFP signal (b). First section (Z1) corresponds to the first Ultrathin section (70nm) where deuterosomes « d » are observed. Deuterosomes are then counted in order of appearance. Sections are selected to have a global vision of all deuterosomes but keep their original number. Scalebar = 2µm. **b**. Cen2GFP signal of the parental centriole-depleted cell observed in EM. « X » indicates Cen2GFP aggregate. Scale bar = 2µm. **c**. Schematic representation of the cell observed in CLEM. Black dots represents deuterosomes, Solid line indicates nucleus. **d**. Serial pictures of a deuterosome from a 0cc cell. Numbers correspond to procentrioles. Scale bar = 200nm. **e**. Correlation between deuterosome diameter and the number of centrioles. Diameter of deuterosomes is calculated on the Ultrathin section where deuterosome surface is the largest. Number of procentrioles is calculated in the same way as in **d.**

**Supplementary Figure 3:**
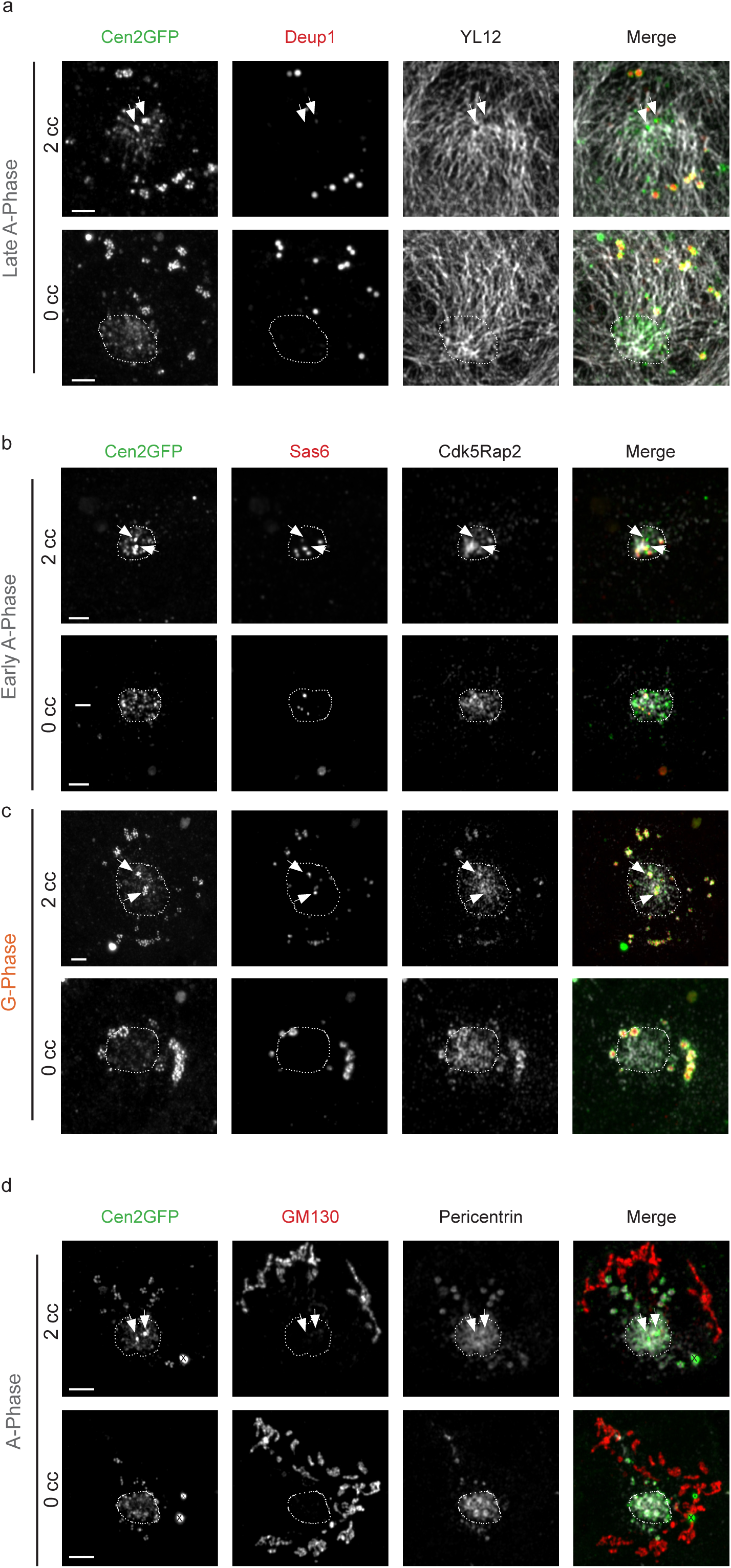
Microenvironment of centriole amplification in centrosomal centriole-depleted cells. **a.** Immunostainings of the tyrosinated microtubule network (YL12) and deuterosomes (Deup1) in 2cc or 0cc cells during Late A-Phase. As for early A-Phase (**Fig 4e**), in 2cc cells, microtubule network converges on one centrosomal centriole whereas in 0cc cells, it converges on the Cen2GFP cloud, where the PCM is enriched. Dashed line delineates Cen2GFP cloud. **b**, **c**. Immunostainings of the PCM protein Cdk5rap2 and Sas6 during A-Phase **(b)** or G-phase **(c)** in 2cc and 0cc Cen2GFP progenitors. Dashed line delineates the PCM cloud. **d**. Immunostainings of the cis-Golgi (GM130) and Pericentrin on 2cc or 0cc Cen2GFP progenitors during A-phase. Dashed line delineates the PCM cloud. Arrows indicate centrosomal centrioles. « X » indicates Cen2GFP aggregate. Scale bar, 2µm.

**Supplementary Figure 4:**
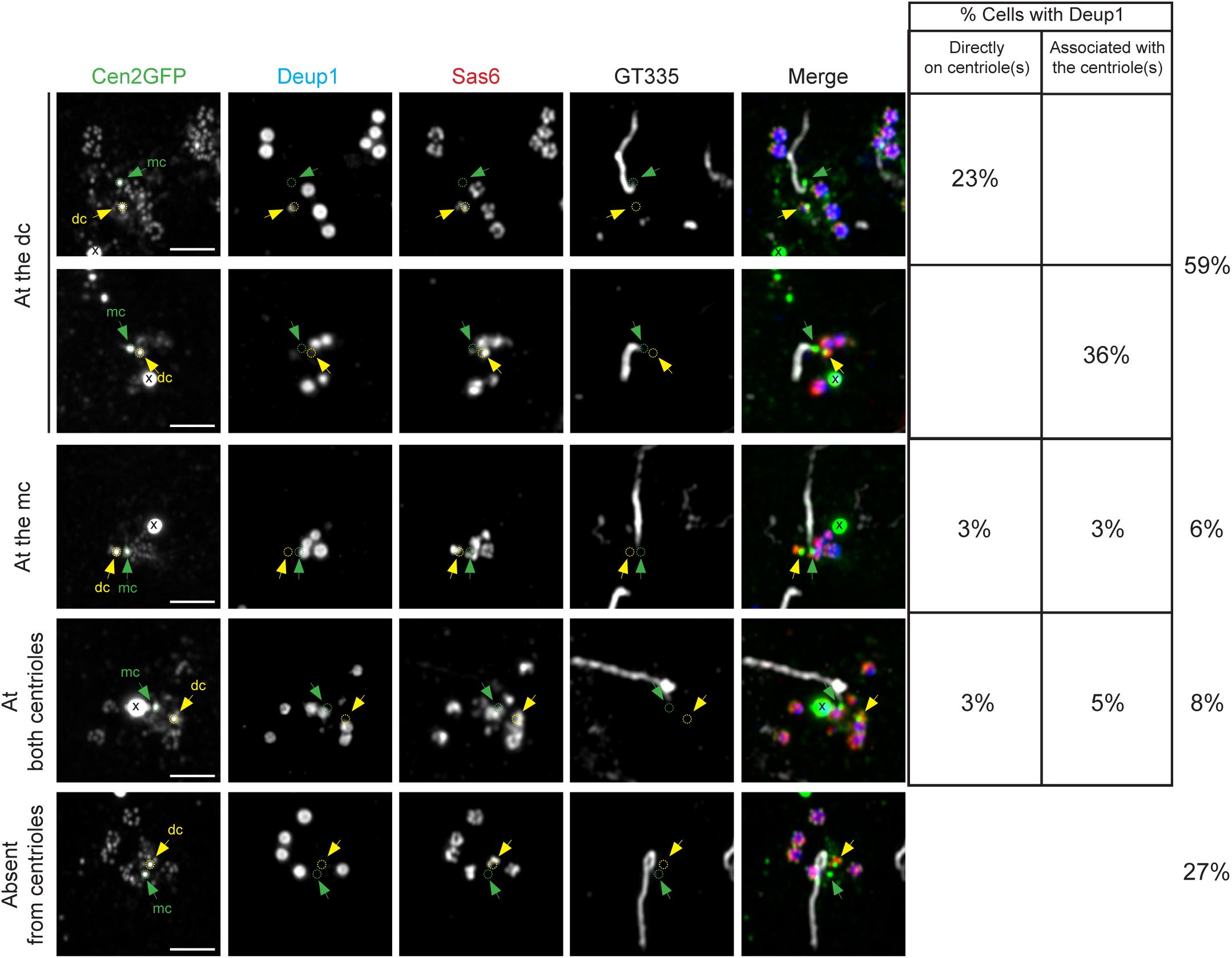
*In vivo* characterization of Deup1 asymmetry on centrosomal centrioles during A-phase. Super-resolution z-projection images and quantification of *in vivo* Cen2GFP ependymal progenitors stained for Deup1, Sas6 and GT335 during A-phase. Association of Deup1 with centrosomal centrioles is assessed by the simultaneous and close presence of Deup1 and Cen2GFP signals on the same z pictures. A-phase identification is based on the Cen2GFP halo signal. Percentages are calculated from 329 cells imaged on the brain lateral ventricular walls from 5 different animals at post-natal day 2 and 4.

**Supplementary Movie 1 Live imaging of centriole amplification in a 2cc Cen2GFP progenitor**

Time-lapse of the centriole amplification dynamic in a 2cc Cen2-GFP progenitor (63X magnification, Δt=40 minutes). « X » indicates Cen2GFP aggregate. Green arrows indicate centrosomal centrioles. Scale Bar, 5 µm.

**Supplementary Movie 2 Live imaging of centriole amplification in a 0cc Cen2GFP progenitor**

Time-lapse of the centriole amplification dynamic in a 0cc Cen2-GFP progenitor (63X magnification, Δt=40 minutes). « X » indicates Cen2GFP aggregate. Scale Bar, 5 µm.

